# Fusome topology and inheritance during insect gametogenesis

**DOI:** 10.1101/2022.05.18.492500

**Authors:** Rocky Diegmiller, Jasmin Imran Alsous, Duojia Li, Yukiko M. Yamashita, Stanislav Y. Shvartsman

## Abstract

From insects to mammals, oocytes ad sperm develop within germline cysts comprising cells connected by intercellular bridges (ICBs). In numerous insects, formation of the cyst is accompanied by growth of the fusome – a membranous that permeates the cyst. Fusome composition and function are best understood in *Drosophila melanogaster*: during oogenesis, the fusome dictates cyst topology and size and facilitates oocyte selection, while during spermatogenesis, the fusome synchronizes the cyst’s response to DNA damage. Despite its myriad and sex-specific roles during insect gametogenesis, fusome growth and inheritance in females and its structure and connectivity in males have remained challenging to investigate. Here, we take advantage of advances in high resolution confocal microscopy and computational image processing tools to reconstruct the topology, growth, and distribution of the fusome in both sexes. Our findings inform a theoretical model for fusome assembly and inheritance during oogenesis, shedding light on symmetry-breaking processes that lead to oocyte selection. In males, we find that cell divisions can deviate from the maximally branched pattern observed in females, leading to greater topological variability. Our work consolidates existing disjoint experimental observations and contributes a readily generalizable computational approach for quantitative studies of gametogenesis within and across species.

**Author summary:** The ubiquity of germline cysts across animals and accelerating advances in microscopy call for quantitative and highly resolved studies of their developmental dynamics. Here we use *Drosophila melanogaster* gametogenesis as a model system, alongside a supervised learning algorithm to study a shared organelle that arises during sperm and oocyte development – the fusome. The fusome is a highly specialized membranous organelle that permeates the cyst in both sexes. Our three-dimensional (3D) reconstructions of the fusome and quantitative measurements at successive stages of cyst development during oogenesis shed light on the evolution of cell fate asymmetry within the germline cyst in females, where the cyst gives rise to a single oocyte. In males, where each cell of the cyst goes on to form sperm, the fusome fragments and exhbits topologies that deviate from the stereotypic maximally branched topology found in females. Our findings can be interpreted in the context of the divergent outcomes of gametogenesis in both sexes and highlight the centrality of quantitative measurements in evaluating hypotheses in biological sciences.

## Introduction

Gamete formation is not a solo act. Across animals, oocytes and sperm develop within germline cysts whose cells are connected by stabilized cytoplasmic channels called intercellular bridges (ICBs) [1–14]. Due to their relatively large size, ICBs enable the intercellular exchange of proteins, mRNAs, and organelles, as well as bulk cytoplasmic flow [6, 8, 15, 16]. Cyst formation and intercellular communication are critical for gametogenesis, enabling cell cycle regulation [17–21], cell fate determination [17, 20–23], as well as sharing of cellular products – even after meiosis [20, 21]. Germline cysts are typically represented as cell lineage trees (CLTs), where cells and ICBs define the nodes and edges of the CLT, respectively [15]; across organisms, CLTs can exhibit substantial differences in size and connectivity patterns [18].

*Drosophila* has emerged as a powerful experimental model for studies of germline cyst formation, cell cycle regulation, and robust cell fate specification [17, 23–26]. In *Drosophila*, sperm and oocytes develop within germline cysts that invariantly comprise 16 cells. To form the cyst, a founder cell – gonialblast in males; cystoblast in females – undergoes several divisions to form a 16-cell cyst, which ultimately gives rise to 64 spermatids in males, and a single oocyte in females (Fig 1A). A prominent feature of the developing germline cyst in *Drosophila* and numerous other insects is the fusome [5, 9, 10, 17, 27–29] – a membranous organelle that grows with each cell division during cyst formation, giving rise to a finger-like backbone that permeates the germline cyst through the ICBs (ring canals in *Drosophila*) [17, 30, 31]. The fusome is rich in vesicular and fibrillar structures and comprises several cytoskeletal proteins such as *α*- and *β*-Spectrin, the adaptor Ankyrin, the adducin-like protein, Hu-li tai shao (Hts), and the spectraplakin Short stop (Shot) [17, 27, 31–34].

**Fig 1.**
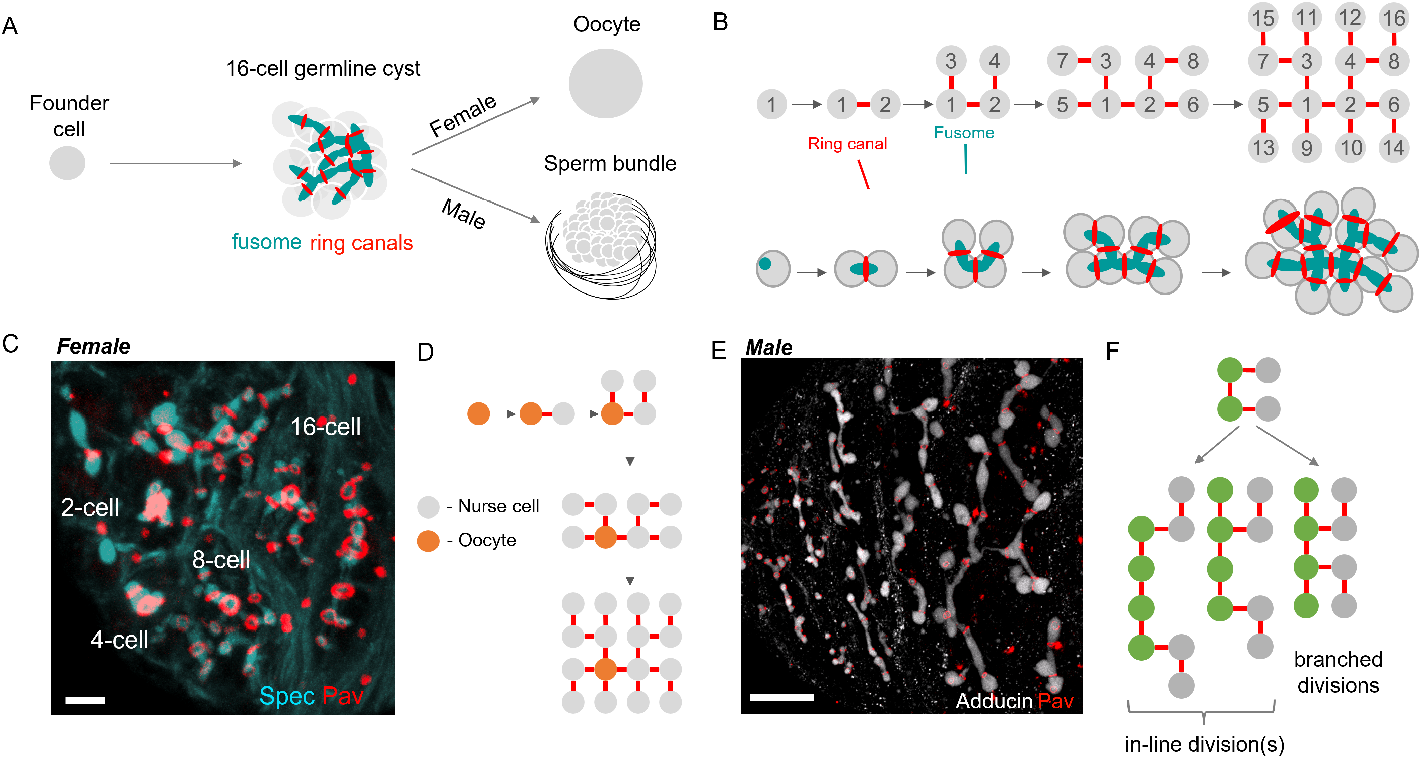
Fusome formation, structure, and patterns of division. A) Schematic of cyst formation during oogenesis and spermatogenesis in *Drosophila melanogaster*. A founder cell gives rise to a germline cyst whose 16 cells are connected through ICBs or ring canals. During oogenesis, one cell becomes the oocyte while the remaining cells serve a supporting role. During spermatogenesis, two subsequent meiotic divisions with incomplete cytokinesis generate 64 haploid spermatids. In both sexes, germline cysts contain a fusome, which permeates the cysts through the ring canals. B) Schematic of the four rounds of synchronous cell divisions that give rise to the female germline cyst, comprising 16 cells (nodes) connected by 15 ring canals (edges). The fusome (cyan) derives from the spectrosome and grows with each cell division; association of the fusome with one of the two mitotic centrosomes orients the planes of cell division, ensuring the stereotypical and maximally branched pattern in female cysts. C) Projection of a 3D image showing the fusome (*α*-Spectrin, Spec) permeating the cyst through the ring canals (Pavarotti, Pav) in cysts of various sizes (2-, 4-,8- and 16-cell cysts, and a Stage 1 egg chamber (EC)). Scale bar = 5 *µ*m. D) Schematic for the ‘pre-determination’ model for oocyte selection, which posits that the initial bias in fusome volume determines which cell becomes the oocyte; as such, oocyte identity is determined at the 1-cell stage. E) Projection of a 3D image of developing cysts during spermatogenesis, where the fusome (Adducin) and ring canals (Pav) are shown. Scale bar = 20 *µ*m. F) Schematic showing that cell divisions can occur in one of two ways based on the orientation that the internal cells (green) divide. If all internal cells divide to form new branches, the cyst will remain maximally branched; however, internal cells can also divide without forming new branches, thus creating in-line divisions and paving the way for toplogies that deviate from the maximally branched pattern.

The fusome plays important but different roles in gametogenesis in males and females. In females, the germline cyst forms a highly invariant and maximally branched 16-cell tree (Fig 1B), which arises from a founding cystoblast that undergoes four rounds of synchronous and incomplete divisions [21, 25, 35]. The fusome arises from a precursor called the spectrosome that is found in the cystoblast [25, 36]. At each cell division in the 2-, 4-, and 8-cell cyst, fusome material is anchored at one end of the mitotic spindle, thus orienting the planes of cell division and ensuring the stereotypical branched pattern of the germline cyst (Fig 1A,C) [17, 34]. The topology of the fusome therefore reflects and dictates the topology of the germline cyst. Notably, intercellular differences in fusome material are implicated in cell fate specification within the female germline cyst. Specifically, asymmetry in fusome inheritance during germline cyst formation is thought to drive the specification of an oocyte through preferential microtubule-dependent transport of oocyte fate determinants to the cell with the greatest amount of fusome [8, 17, 19, 21, 37–39]. However, despite the fusome’s central role during oogenesis, the dynamics of fusome growth and its inheritance within the growing germline cyst have not been quantified.

In males, where all cells are equivalent and give rise to 64 spermatids through two additional meiotic divisions [4, 19, 21], the fusome also plays a part [21, 40–42]. Here, the fusome is thought to be dispensable under normal conditions [19]; however, in cases of DNA damage, the fusome facilitates synchronization of spermatogonial death, thereby shielding the integrity of gamete genomes [20]. Formation and growth of the fusome during spermatogenesis is thought to occur through a similar sequence of divisions as in oogenesis, where synchronous divisions of a cystoblast are accompanied by concomitant growth of the fusome, leading to a maximally branched CLT. However, fusome topology within male cysts and its effects on cyst topology are less understood than in females and have yet to be confirmed through direct reconstructions.

To address these gaps in knowledge, we leveraged advances in high resolution confocal microscopy and a recently developed and tested pipeline for 3D image analysis and quantification [43] to measure and reconstruct the fusome and its growth dynamics at all stages of cyst formation in both sexes (Fig 2). In females, we show that cystoblast divisions are not inherently asymmetric with respect to fusome partitioning and that, instead, newly generated fusome fragments are shared equally between mother and daughter at each division – as predicted by a minimal mathematical model of additive fusome growth. In males, we demonstrate a greater diversity of germline cyst topology than previously appreciated, and reconcile these findings with the divergent functions of the fusome in both sexes. The algorithm developed here is readily extended to any experimental system comprising cells connected via ICBs, such as those that arise during animal gametogenesis [1–13]. Our work thus contributes a valuable computational tool that can recover key morphological features of germline cyst development, thus permitting comparative studies of gametogenesis within and across species.

**Fig 2.**
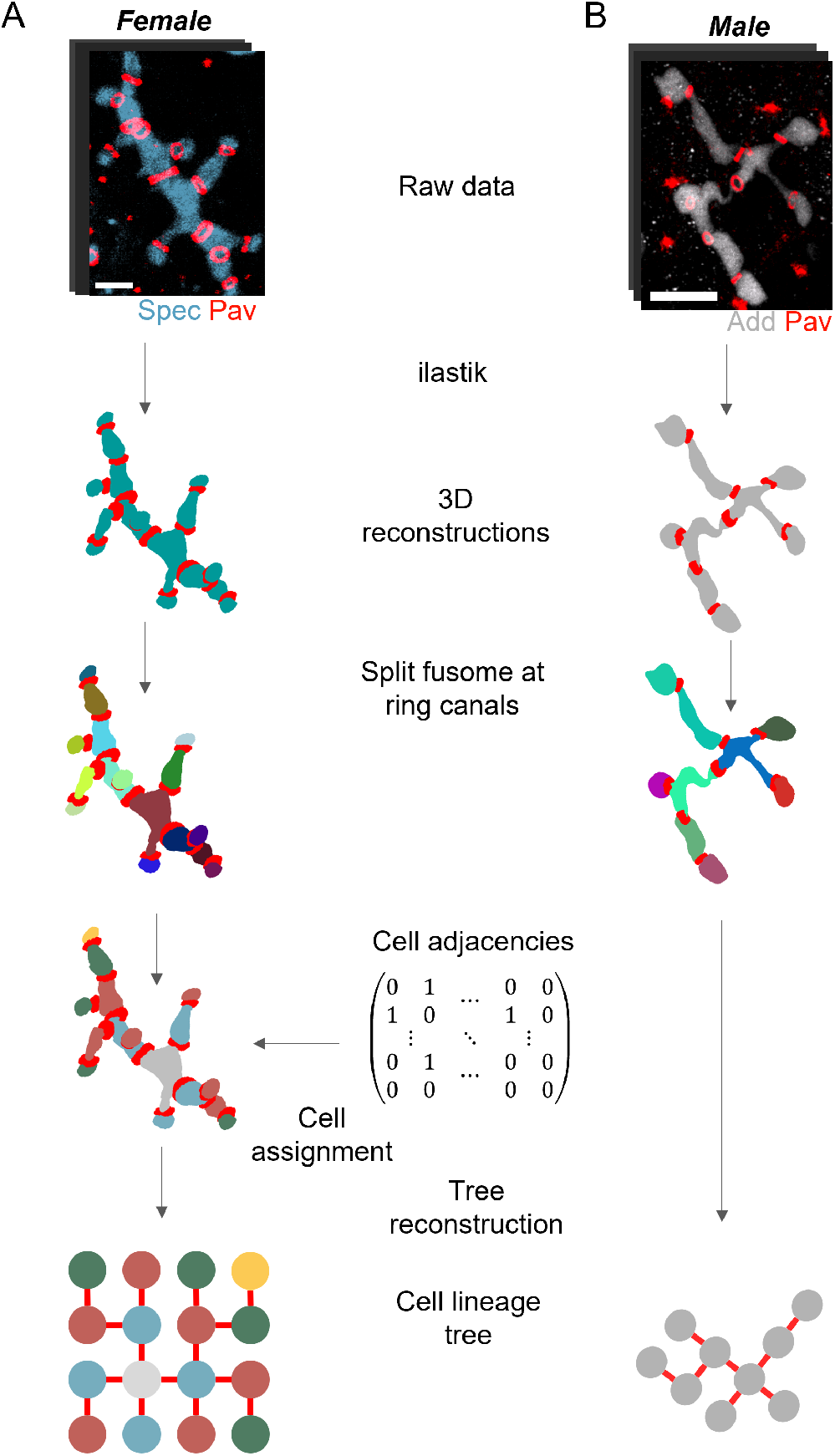
Pipeline for 3D fusome reconstructions in developing germline cysts. (A) Starting from 3D confocal images of fusomes with fluorescently-labeled ring canals and fusome, training in a machine learning program and processing in MATLAB as described in [43, 44] produces a 3D reconstruction of the fusome and ring canal system at successive stages of germline cyst formation (Figure 1A). From here, ring canals at the junction between sister cells are used to segment the fusome into its component parts (multicolored cells) and, in the case of oogenesis, produce an adjacency matrix representing intercellular connections within the cyst to compare with the known female germline cyst network topology, permitting unambiguous cell label assignments to the individual fusome fragments. These fusome fragments are then labeled according to their position relative to the presumptive oocyte. (B) Male cysts underwent the same procedure of collective adjacency information, but as no oocyte is present, no distinction was made between cells. Scale bars = 2 *µ*m (A) and 10 *µ*m (B).

## Results

### Supervised learning for network topology reconstruction

To analyze fusome topology and intercellular volume distribution during cyst formation in females, we implemented a supervised learning algorithm to identify the location of fluorescently-labeled fusome and ring canals within the developing cyst (Fig 2 – as previously described [43, 44]. Briefly, for each cyst, the fusome (fluorescently-labeled through tagging of *α*-Spectrin in females and adducin in males) and the ring canals (through endogenously-tagged Pavarotti-GFP, a key component of ring canals in *Drosophila*) were identified using the supervised learning program ilastik [44, 45], which has been used in several other settings for volumetric reconstructions of cells and subcellular structures [43, 46–48]. The fusome in females was taken as a single, simply-connected component within the developing germline cyst. Ring canals did not have this constraint, and based on the sample under consideration, either 1, 3, 7, or 15 objects were required to complete the fusome segmentation and reconstruction upon using skeletonization to identify objects which were closed loops within each image stack (corresponding to 2-, 4-, 8-, and 16-cell cysts, respectively). Each ring canal, which necessarily surrounds the fusome at cell junctions, was used as a marker to splice the fusome into multiple sections. A recursive function was then built so that for each distinct ring canal, the fusome was split into two fragments on either side.

At the first iteration, the fusome was split into two fragments. The algorithm then iteratively split the fusome at each ring canal, generating fragments that corresponded to the volumes present within each cell of the cyst. After generating these fragments, the function then logged information about their volumes and positions within the stack and moved to a different branch of the recursive loop. The final output of this algorithm was a three-dimensional stack of separated fusome fragments, whose sum amassed the entirety of the original fusome. These objects were then exported for further identification based on mutual adjacencies to assign a unique label to each piece of fusome based on its number of neighbors and distance from the presumed founder cell (in females). The volume of each fragment was calculated based on the number of voxels within each object, multiplied by the known voxel size.

We found that our pipeline reconstructs the correct maximally branched cyst structure in females (Fig 2A). Because the topology of the network at each division is symmetric during oogenesis, the cell with the larger fusome fragment and four ring canals was labeled cell 1. Each cell thereafter was uniquely identified based on its number of neighbors and proximity (number of intervening ring canals) to cell 1.

From here, the volumes of fusome within each cell in the cyst were also recorded to enable intercellular comparisons in the dividing cyst. Manual validation of ilastik volumes was performed on a subset of randomly selected samples. For each sample, manual thresholding was performed in FIJI to identify fusome area at each slice. All slices were summed and the total was multiplied was the voxel size to yield a fusome volume estimate. Across all samples, the difference between manual and automated fusome volumes was*∼* 8% on average (File S1).

To further test and extend the applicability of the developed algorithm, we turned our attention to the male, where we analyzed the 3D structure and topology of fusomes at successive stages of cyst formation (Fig 2B) [43, 44]. In contrast to the female cyst, where cells were tightly packed and the fusome compact, male germline cysts appeared more elongated, with thread-like fusomal materials connecting otherwise seemingly disjoint fusome fragments, as highlighted previously [19, 37, 42, 49].

### Quantifying fusome growth dynamics during oogenesis

Intercellular asymmetry in fusome distribution is thought to mediate oocyte fate specification and polarity, which in *Drosophila*, ultimately establish the embryo’s main body axes [17, 25, 26]. According to this ‘predetermination’ model for oocyte selection, the cell that inherits the greatest amount of fusome becomes the oocyte, and that cell is always one of the two cells with four ring canals – the pro-oocytes (Figure 1C). This model builds on the fact that asymmetry in fusome content arises during the very first cell division, and that this asymmetry between the first cell and remaining cyst cells, including the other pro-oocyte, is maintained during subsequent divisions [27, 36, 50]. As microtubules associate with the fusome, its asymmetry is proposed to predetermine oocyte identity by directing the microtubule network towards one cell, along which dynein transports oocyte fate determinants [25, 30, 51, 52]. Formation of a polarized microtubule network is indeed key for oocyte specification [23, 52]. For example, *egalitarian*, in concert with *Bicaudal-D*, promotes the microtubule-dependent transport of factors to the presumptive oocyte; its mutants form cysts with 16 nurse cells despite forming normal fusomes that appear asymmetrically distributed [53–56]. The alternative model for oocyte selection postulates that fusome asymmetry is insufficient for determining oocyte fate; instead, the pro-oocytes are considered equivalent, with both having an equal chance of becoming the oocyte [25, 57]. According to this model, one of the pro-oocytes becomes the oocyte ‘stochastically’, through competition for some limiting oocyte determining factor(s) at the 16-cell stage (Figure 1C).

Quantification of fusome growth, inheritance, and relative fusome volumes within connected cells is key for distinguishing between models; however, these measurements are technically challenging. A further complication is the fact that oocyte identity only becomes clear once the 16-cell cyst of *Drosophila* is formed – a point at which several fusome components have begun to disintegrate [25, 30, 57, 58]. Using the supervised learning approach described above, our data revealed a strong bias in fusome volumes between the pro-oocytes starting at the 2-cell stage. We found that the larger of the two fusome fragments comprises, on average, 70% of the total fusome volume. Subsequent divisions however decreased the asymmetry between the two most central cells, with the ratio of volumes between cells 1 and 2 decreasing from 2.35 *±* 0.29 in the 2-cell cyst (*n* = 12) to 1.25*±* 0.18 in the 16-cell cyst (*n* = 11) (Fig 3). Nonetheless, at each stage, there was a persistent fusome volume bias toward these two central cells compared to the other 14: upon successive divisions to form a 4-, 8-, and finally 16-cell cyst, the volume fraction of these two cells was 77.5*±* 6.7% (*n* = 12), 58.2 *±*5.7% (*n* = 6), and 37.1 *±*3.0% (*n* = 11), respectively, dwarfing the fusome portions in each other cell at every stage of division.

**Fig 3.**
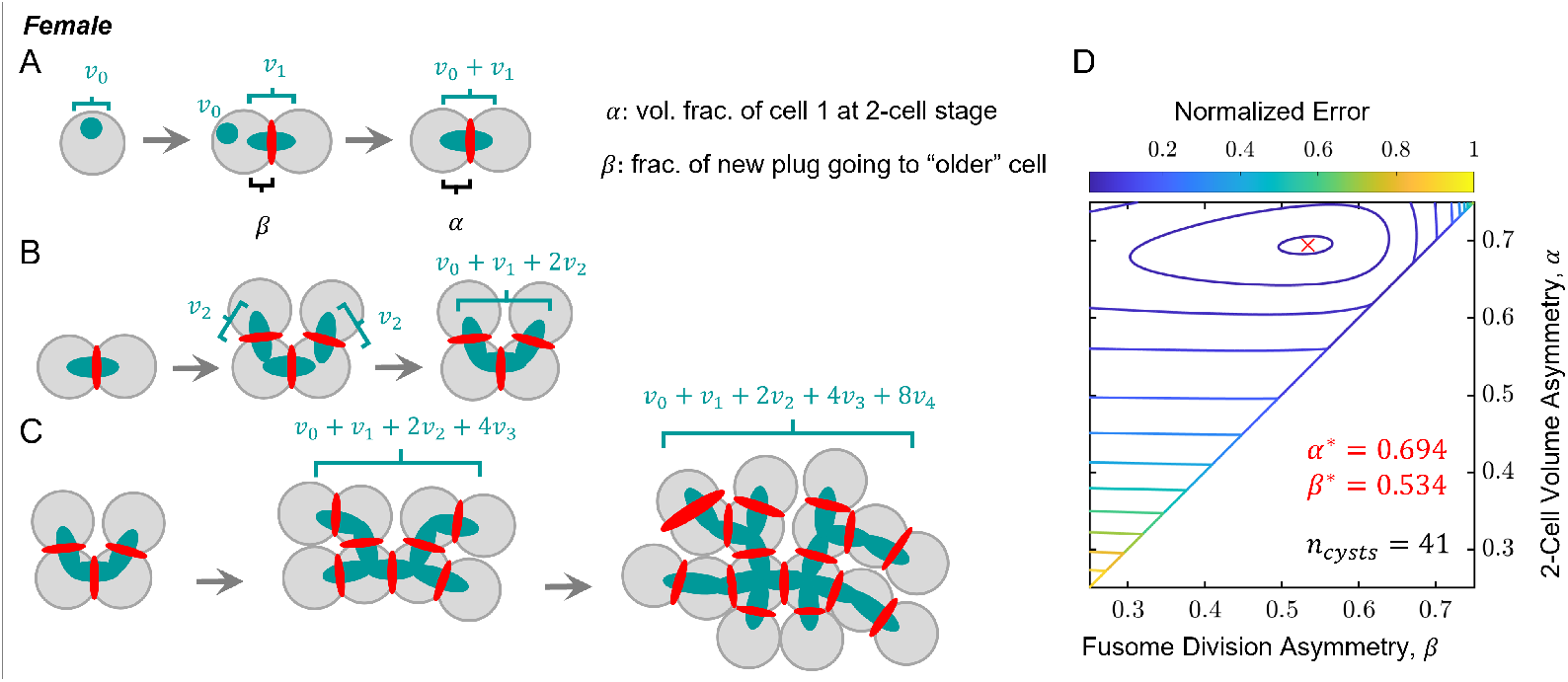
A model for fusome growth and inheritance during oogenesis. A) Starting with a spectrosome of volume *v*_0_ and fusome plug at the first division of volume *v*_1_, the volume fraction *f*_1,2_ of the 2-cell female cyst can be represented as shown in Equation 1, where the volume fraction in cell 1 at this stage is *α* and the asymmetry in sharing of the newly formed fusome component between mother and daughter cell is *β*. B) At the 2-to-4-cell transition, two new fusome plugs are added in the same manner as in A, with each contributing volume *v*_2_. C) Divisions occur again to form 8- and 16-cell cysts, where the total fusome volume can be tracked throughout cyst formation, and expressions for volume fractions of fusome within each cell at each cyst size to be derived. D) Error calculations for ranges of the model parameters *α* and *β*. For each set of parameters, model volume fractions for each cyst size were calculated and compared with measured data by summing over the squared residuals between measured and derived volume fractions over each cell at all cyst sizes. The minimum error using this metric was found at the value *α* = 0.694, *β* = 0.534, corresponding to a scenario where fusome plugs are added evenly between the mother and daughter cells at each successive division. Explanation of model assumptions and equations can be found in the Materials and Methods. Comparison with measured data can be found in Fig S1.

We also found that aggregating the volume fractions based on ring canal number provided a meaningful metric with respect to when each cell is formed in the successive divisions. In particular, we found that cells with 4, 3, 2, and 1 ring canals comprised 37.1 *±*3.0%, 21.4*±* 2.6%, 21.7 *±*2.4% and 19.8 *±*2.6% of the total fusome volume(*n* = 11), respectively. Note that the two cells with 4 ring canals are the pro-oocytes. Within the measurement error of each group and for comparison with theory, these groupwise fractions at the 16-cell stage are taken as the following ratio: 2 : 1 : 1 : 1.

To rationalize these experimental data, we developed a minimal theoretical framework to model how relative fusome volume fraction in each cell changes during cyst generation. This model is built on the fact that the fusome in *Drosophila* grows through successive rounds of accumulation and fusion of fusome fragments in the ring canals connecting mother and daughter cells [25, 50, 57, 58]. Here, we only assumed that at each division there was no additional change in the relative volume fractions of fusome in the cells between divisions, other than what was added from the fusome plug [25, 50]. This model therefore treats the fusome as an irreducible building block, where plugs chain together at each division to form a larger and branched structure.

Building on these assumptions, we wrote a series of equations with two parameters: *α*, the volume fraction of fusome in the founder cell at the 2-cell stage, and *β*, the fraction of the fusome plug added to the parent cell at each division (see Materials and Methods). Using the 2 : 1 : 1 : 1 heuristic for the groupwise fractions at the 16-cell stage and inserting the derived relationships into equations for the volume fractions for each cell in the 2-, 4-, 8-, and 16-cell cysts yields volume fractions that enable direct comparison of model predictions with experimental measurements over a range of values for *α* and *β* (Fig 3D). For the parameter range *α∈* [0.25, 0.75] and *β∈* [0.25, 0.75], we found *α* = 0.694 and *β* = 0.534 minimized the error with our measurements, yielding for a range of values for *α* and *β*. Considering the range of parameters that yield errors within 5% of the minimum gives ranges of *α∈* (0.683, 0.705) and *β ∈* (0.497, 0.568).Notably, this interval for *β* captures the value 0.5, suggesting that fusome fragments are shared equally upon division [25, 50].

### Reconstructing fusome and cyst topology during spermatogenesis

In any interconnected cyst, two types of cells exist: terminal cells – the cells at the very ends of the cyst, connected to only one other cell – and internal cells. Upon division, terminal cells form one cell connected to two cells and another cell that now again has only one connection (Fig 1F). In other words, terminal cell divisions results in an internal cell and a terminal cell. On the other hand, internal cells can divide in two ways: they can forming a new terminal cell or two internal cells. These division types affect CLT topology: maximally branched trees form when all internal cells divide to form terminal cells (Fig 1F); however, if in-line divisions occur, maximal branching is lost. By tracking cell-cell connections, the topology of the germline CLT can be extracted [59, 60]. With these division rules in mind, we set out to catalog the series of connections present in male germline cysts and to compare those to the known sequencem of divisions during *Drosophila* oogenesis.

Reconstructions of fusomes in developing male cysts revealed significant fusome fragmentation (Fig 4A). Indeed, we found that in only (50%) of reconstructed samples was the number of fusome-connected cells 2^*n*^, in contrast to females, where in all samples, the number of fusome-connected cell was 2^*n*^. This deviation in males is unlikely to be due to an atypical number of cells. Indeed, when we counted the spermatid tails per sperm tail bundle present across a number of samples, we observed an average of 63.15*±* 1.31 (*n* = 61) spermatids per bundle in post-meiotic cysts, consistent with each cyst comprising 16 cells (Figure S1) [61, 62]. To explore the effects of fusome fragmentation, we analyzed the various topologies that could be recapitulated by tracing fusomal connections in males, assuming that fusome topology reflects that of the CLT. Starting from a single cell, there is only one path towards forming a four-cell cyst via synchronous divisions: a linear fusome fragment with three ICBs. Upon dividing synchronously again, multiple eight-cell cysts are possible, depending on whether the divisions occur in a maximally branched fashion; that is, whether or not each division forms a new terminal fusome fragment budded off from the parent (Fig 1F). For cells at the terminal ends of the cyst, the fusome can only elongate in one manner; however, for the inner cells, the fusome either adds a branch or continues growth in a linear manner. By symmetry, one is left with three possible configurations for the cyst; of these, only one configuration is maximally branched, whereby each division creates a new branch (Fig 1F). Of all fusome reconstructions encompassing 5-8 cells (n = 21), we found none that were inconsistent with the maximally branched connections of an 8-cell cyst (Fig 4A).

**Fig 4.**
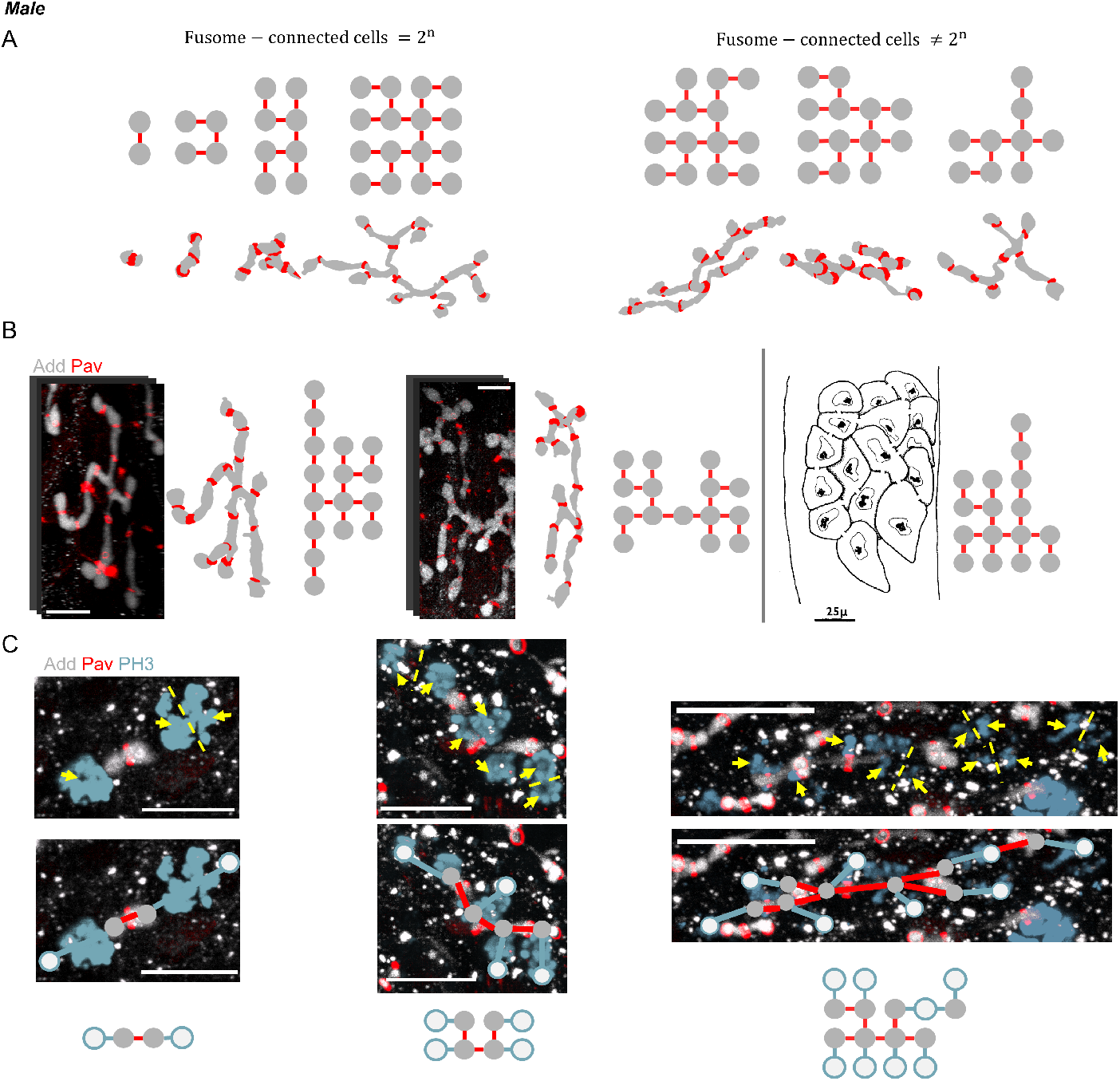
Fusome fragmentation during spermatogenesis and loss of maximal branching. A) Starting with *n* = 62 reconstructed fusomes, a range of cyst sizes can be inferred by identifying the largest connected portions of fusome within each sample. These fusome portions, and their corresponding ring canals, can be used to identify the number of cells connected by each fusome fragment (fusome-connected cells). In approximately half of these, the number of fusome-connected cells was a power of 2 (n=31). B) *Left:* Two examples of fusomes connecting 16 and 15 cells, respectively, whose topology cannot be embedded onto that of a maximally branched network. *Right:* Hand-drawn reconstruction of the cells and intercellular connections of a 16-cell sperm cyst from Figure 1 of the 1973 work by Rasmussen ([41], reproduced with permission). C) Projections of 3D images of dividing cysts with labeled fusome (Adducin, gray), ring canals (Pav, red), and mitotic chromosomes (Phosphohistone 3, cyan), showing examples of a 2-cell (left), 4-cell (middle), and 8-cell cyst (right) admist mitosis. In at least one location in the 8-cell cyst, cells are dividing without forming a new branch; as such, the resulting 16-cell tree will not possess a maximally-branch topology. In all examples, yellow dotted lines denote the likely plane of division for actively dividing cells, while yellow arrows depict clear bundles of aggregated or migrating chromosomes. In addition, nodes representing the locations of existing cells, as well as the likely positions of newly-formed cell based on the plane of division are shown. Scale bars = 10 *µ*m.

Starting with maximally branched 8-cell cysts then, 36 possible configurations are obtainable through another division to form a 16-cell cyst, with only one resulting in a maximally branched network. We analyzed fusome reconstructions connecting 9-16 cells and, while we observed fusomes whose structure was consistent with a 16-cell maximally branched cyst, we uncovered fusomes whose connections could not be embedded onto a maximally branched topology. Even in cases where fusomes did not connect 16 cells, the topology of the largest fragment was sufficient to demonstrate that divisions had not occurred in a maximally branched manner (Fig 4B). We next analyzed the orientation of mitotic spindles in dividing cysts, focusing on the fourth division cycle, when the cyst increases in cell number from 8 to 16. In female cysts, mitotic spindles align such that newly-formed cells divide to create a maximally branched tree as opposed to ‘in-line’ (Fig 1B); the fusome follows cyst growth, leading to its branching throughout the cyst [17]. This division pattern was observed in all dividing 2- and 4-male cysts, but not in 8-cell cysts, where we found mitotic spindles that were parallel to two disjoint fusomal fragments (Fig 4C) – consistent with cell division elongating the fusome rather than giving rise to a new branch. These findings suggest that the stereotyped pattern of connections in female cysts are not strictly preserved during spermatogenesis and that deviations from maximally branched structures in male cysts arise at the fourth division during cyst formation.

## Discussion

The fusome has long been identified as a prominent feature of developing germline cysts in insects; first described in spermatocytes more than a century ago, the fusome has since been observed in numerous species [5, 9, 10, 17, 27–29]. Fusome composition and function are best described in *Drosophila* [17, 25, 27, 30, 32, 35, 36, 50], yet questions remain regarding its growth and inheritance pattern in females, as well as its topology in males. Here we set out to address these gaps in knowledge by adapting a machine-learning approach for 3D reconstructions of cells and their substructures [43]. In females, the nonuniform distribution of the fusome has been implicated in oocyte selection [17, 30]. The ‘predetermination’ model for oocyte selection posits that the slight asymmetry provided by the fusome biases the choice of oocyte to the first cell – a fate that is then sealed by directing flow of other determinants [34–36, 50]; the alternative ‘stochastic’ model for oocyte selection suggests that pro-oocytes are equivalent, but does not account for how oocyte selection is restricted to those two cells [24, 57, 63].

Based on direct measurements of fusome volumes in daughter cells at successive stages of germline cyst formation, we show that cystoblast divisions are not asymmetric with respect to fusome partitioning; instead, our data are consistent with a model in which newly generated fusome fragments are shared equally between mother and daughter at each division and where the fusome is assembled in an additive manner. Therefore, asymmetry in fusome distribution within the cyst arises by propagating the advantage that the founding cystoblast has by virtue of inheriting the spectrosome. Consequently, fusome asymmetry between the two pro-oocytes decreases with each cell division, while asymmetry between the pro-oocytes and the rest of the cyst increases as the volume fraction shared between these two cells dwarfs that in all other cells in the cyst.

Based on these data, the strict dichotomy between the two oocyte selection models is likely to be overstated; further, neither model is consistent with all previous experimental observations of mutants in which oocyte selection is inhibited [57], delayed [64], or where multiple oocytes are specified [65, 66]. Instead, we propose a third model of ‘equivalency with a bias’, whereby the amplified fusome asymmetry between the pro-oocytes and the remaining cells biases the choice of oocyte to the two pro-oocytes, while the decreasing fusome asymmetry between the two pro-oocytes themselves means that which of the pro-oocytes is chosen as the oocyte at the 16-cell stage is likely to be stochastic. Such a two-step mechanism, one that relies on the interplay of predetermined factors to limit the pool of candidates and stochastic factors for the final selection has been documented in other well-studied systems of robust cell fate determination, such as the invariant spatial patterning of the equipotent vulval precursor cells in C. elegans and the achaete(ac)-scute(sc) system in bristle determination in the adult *Drosophila* cuticle [67–70].

In males, the fusome is also a prominent structure, and while it is thought to be unimportant for normal development [19], it appears to play a crucial role in mediating intercellular connectivity and synchronized cell behaviours under stress [20]. Male fusomes also last longer: while fusomes start to disintegrate following formation of the 16-cell cyst in females, male fusomes persist following meiosis [39]. Despite these inherent differences, the topology of the male fusome, and consequently that of the germline cyst, have long been assumed to mirror those of female maximally branched structures. By analyzing fusomes at successive stages of cyst development, we instead found that division orientation in male cysts exhibits greater variability those in females, particularly where the fusome has become thread-like or fragmented.

To our knowledge, the reconstructions of the fusome-ring system within the male germline cyst presented here are novel; however, we are not the first to demonstrate that male cysts may exhibit ‘atypical’ division patterns: a manual reconstruction by Rasmussen fifty years ago demonstrated that the structure of cell-cell connections did not align with the known maximally-branxhed pattern found in female cysts [41] (Fig 4B). The reconstruction and observations made my Rasmussen have since been largely lost to the field, with some authors later stating that “the distribution of ring canals among the cells of the cyst appears to be the same as described by King for oogenesis…; the recent report to the contrary by Rasmussen probably represents an exceptional cyst.” [71].

Rasmussen concluded that the observed atypical male cyst must have formed as the result of five mitotic divisions [41]. Alternatively, as our results suggest here, cysts without a maximally branched structure can arise through four synchronous mitotic divisions, where the final division occurs without maximal branching. Consequently, and in contrast to females, we find that the number of ring canals in males does not strictly reflect the history of divisions within the cyst. These findings can be interpreted in the context of the divergent outcomes of gametogenesis in both sexes: a cyst whose cells all share the same fate in males, and a cyst that produces a single oocyte in females through the directed transport of materials by the other 15 supporting nurse cells. Compared to females, males may therefore have less stringent requirements for setting up a polarized microtubule transport network that would require an intact fusomal backbone that is asymmetrically partitioned within the cyst.

Reconstructions of germline cysts in various species have revealed a slew of diverse topologies, from linear chains in the plumed worm to star-like cysts in the white worm where cells connect to a central cytophore [1, 18, 72, 73]. Recent studies have also revealed the presence of ICBs in choanoflagelates – a model organism for studies of the origins of multicellularity [43, 74]. While the results here are derived based on the known division patterns during *Drosophila* oogenesis, the developed analysis pipeline is readily generalizable to any species whose germline cysts develop with the aide of fusomal material, such as those of the order Lepidoptera [75, 76] and those that do not, like the mouse [77]. The approach presented here will therefore be key for expanding the zoology of cyst structures in the animal kingdom and for investigating the interactions between cyst structure and size, and how these vary within and across species [18, 78, 79].

## Materials and Methods

### Fly stocks and sample preparation

Pav-mCherry,orb^*F* 343^ females (gift from Schedl lab, Princeton University) were used in this study as a marker for ring canals and to allow for isolation of 16-cell cysts. Ovaries were dissected and fixed as previously described, taking care to isolate germaria from ovarioles [43, 80]. After blocking, the primary antibody mouse anti-*α*-Spectrin 3A9 (1:30, gift from Schedl lab, Princeton University) was added and rocked overnight at 4^*◦*^C overnight. The secondary antibody rat anti-mouse Alexa-Fluor 488 (1:100, Invitrogen) was then added to mark the fusome within each 2-, 4-, 8-, and 16-cell cyst in the germaria. Dissected ovaries were mounted in a 75:25 mixture of RapiClear 1.47 (SUNJin Lab) and Aqua-Poly/Mount (Polysciences).

Ubi-Pavarotti-GFP male flies (gift from Glover lab, California Institute of Technology) were dissected according to standard protocols [45]. Immunofluorescence staining was performed as described previously [81]. Briefly, tissues were dissected in PBS, transferred to 4% formaldehyde in PBS and fixed for 30 minutes. Tissues were then washed in PBS-T (PBS containing 0.1% Triton X-100) for at least three 10 minute washes, followed by incubation with primary antibody in 3% bovine serum albumin (BSA) in PBS-T at 4^*◦*^C overnight. Samples were then washed for three 20 minutes washes in PBS-T, incubated with secondary antibody in 3% BSA in PBS-T at 4^*◦*^C overnight, washed as above, and mounted in Vectashield with DAPI (Vector Labs). The following primary antibodies were used: rabbit anti-Phospho H3 (Ser10) (1:200, Thermo Fisher) and mouse anti-Hts (1:20, 1B1, Developmental Studies Hybridoma Bank). Secondary antibodies used were anti-rabbit IgG Alexa Fluor 488 and anti-mouse IgG Alexa Fluor 647 (both 1:200). Fluorescent images were acquired using a Leica TCS SP5 confocal microscope with a 63x/1.3 NA oil objective.

Confocal imaging was performed on a Leica SP5 confocal microscope using a 63x/1.3 NA oil objective. Three dimensional stacks were acquired using 488 and 546 nm lasers in series between roughly 100 slices of 16-bit, 1024 × 1024 images. Acquired images were roughly 70 nm in x and y, and 210 nm in z.

### Spermatid sample preparation and imaging

Testes were fixed for 1 h or overnight (at 4°C) with 2.5% glutaraldehyde in 0.1M Sorensen’s buffer, pH 7.4. Samples were rinsed twice for 5 min each with 0.1 M Sorensen’s buffer and postfixed for 1 h in 1% osmium tetroxide in 0.1 M Sorensen’s buffer. Next, testes were rinsed twice in double distilled water for 5 min each and en bloc stained with 2% uranyl acetate in double distilled water for 1 h. The samples were then dehydrated in increasing concentrations of ethanol, rinsed with acetone, and embedded in epon epoxy resin. Thin sections were mounted on Formvar/carbon-coated slotted grids and poststained with uranyl acetate and lead citrate. Samples were examined on a JEOL1400 transmission electron microscope and images captured using a sCMOS XR401 custom-engineered optic camera by AMT (Advanced Microscopy Techniques).

### Image acquisition and processing

Raw image stacks were first preprocessed to isolate individual cysts within each germarium. Stacks were uploaded into ilastik for Pixel Classification to identify the locations of fusome and ring canals for each sample, in a manner similar to previous work [43, 44]. For each type of data, training was performed by manually identifying pixels that belong to each class of interest. This positional information, a set of probabilities at each voxel for belonging to fusome, ring, or image background, were imported into MATLAB for object identification and volume measurement. Using the probabilities for each voxel to be part of the fusome and ring canals, simple thresholding was used to find the most likely positions for each class.

Due to the invariance of cyst structure in the *Drosophila* female germline, the adjacencies between cells in the female cyst are known, up to identification of one cell due to cyst symmetry. In this case, larger fusome fragment with the most ring canals were taken to be the founder cell (cell 1), making it possible to assign all other identities within each cyst. The adjacencies for each set of fusome fragments were identified and mapped to the known adjacency matrices at each cyst size, allowing for reassignment through the Hungarian algorithm as previously described [43, 82].

In males, fusome fragmentation required for manual segmentation or validation to ensure proper cell-cell adjacencies were collected. Samples stained with Phosphohistone 3 were additionally trained within ilastik and inserted into MATLAB for further analysis for positioning. 3D reconstructions allowed for the identification of cells dividing without the formation of new fusome branches.

### Mathematical model for fusome assembly in females

In this section, we introduce a theoretical model for fusome formation during *Drosophila* oogenesis using known biological features of the process, accompanied by a small number of assumptions. This minimal model, in coordination with the measured fusome volumes across stages, seeks a more coarse-grained, less stochastic view of fusome formation during oogenesis.

It was previously shown that the cystoblast contains the spectrosome, a fusomal precursor [25, 36]. Therefore, if the volume of the spectrosome in the cystoblast after division to form the 2-cell cyst is given by *v*_0_ and the fusome volume added at the first division is *v*_1_, the volume fraction of cell 1 in the 2-cell cyst (*f*_1,2_) can be described by:

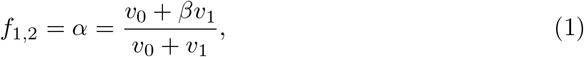

where *α* is defined as the volume fraction of fusome in cell 1 after the first division and *β* is defined as the fraction of the fusome volume added to the already existing cell at each division (Fig 3A). Extending this quantitative description to the next division, from a 2-cell to 4-cell cyst, we have two fusome plugs to add, each contributing *v*_2_ to the total fusome volume (Fig 3B). Under the assumption that fusome plugs added at each division are the same (in this case, *v*_2_), the volume fractions in the cells of the cyst, *f*_*i*,4_, can be derived:

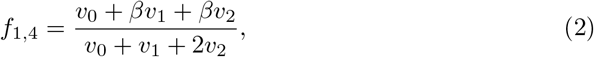

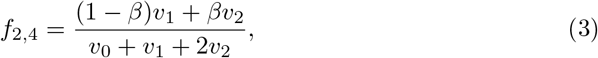

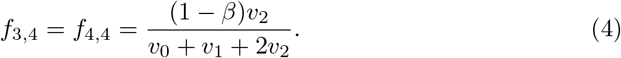

These derivations can be repeated at each subsequent division, allowing for fusome volume fractions in each cell of any cyst size to be defined (Fig 3A-C).

Our experimental measurements revealed that the average volume fractions of cells 1 and 2, cells 3 and 4, cells 5 through 8, and cells 9 through 16 at the 16-cell stage are given in the ratio 2 : 1 : 1 : 1. One can therefore write the following equations for the volume fractions of these groups of cells and, along with Equation 1, solve for *v*_1_, *v*_2_, *v*_3_, and *v*_4_, the average volume of the fusome fragment being added between mother and daughter at each division, in terms of *v*_0_:

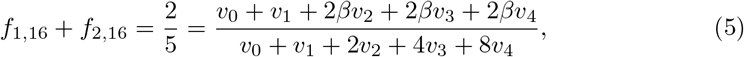

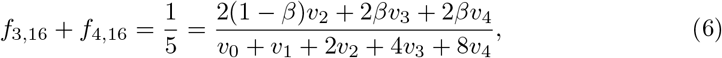

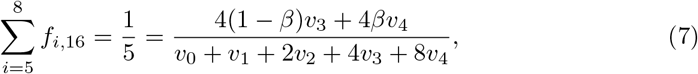

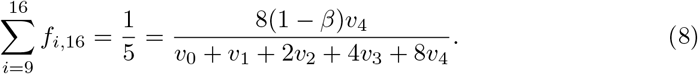

Solving this system of equations, along with 1, allows for one to solve for *v*_1_, *v*_2_, *v*_3_, and *v*_4_ in terms of *v*_0_. Inserting the derived relationships into equations for the volume fractions for each cell in the 2-, 4-, 8-, and 16-cell cysts yields volume fractions that can be compared with experimental measurements at the parameter values *α* = 0.7 and *β* = 0.5 that were found to minimize the errors, as shown above (Fig 3D, Fig S1).

### Data availability

All codes used in the reconstruction and analysis of these data can be found at https://github.com/Shvartsman-Lab/FusomeFormation or https://github.com/Shvartsman-Lab/MaleReconstruction. Sample input image stacks, ilastik training sets and output files for these codes will be made available upon request.

## Supporting information

**S1 Fig. Comparison of female fusome volume fraction model with experimental data**. Plot of measured average volume fractions for 2-cell (*n* = 12), cell (*n* = 12), 8-cell (*n* = 6), and 16-cell (*n* = 11) cysts (black circles) compared with theoretically predicted values (orange circles) for *α* = 0.7, *β* = 0.5. Adjacent to each plot are color-coded schematics of the cyst networks and representative reconstruction of the fusome. Errors bars represent standard deviation.

**S1 File. Female fusome volumes and validation by manual reconstructions**. Fusome volumes and fusome volume fractions are shown from measured automated reconstructions from ilastik and MATLAB. The list comprises measured volumes from 2-cell (*n* = 12), 4-cell (*n* = 12), 8-cell (*n* = 6), and 16-cell (*n* = 11) cysts, along with estimated volume fractions at each stage given the estimate of *α* = 0.7, *β* = 0.5 for the values of the model parameters. In addition, manual volume reconstructions on a randomly chosen group of samples (*n* = 8) using thresholding in FIJI shows a high agreement between manual and automated fusome volume reconstruction in 3D.

**S2 File. Spermatid tail counts**. Spermatid tail bundles were cut and visualized in FIJI. Manual counting across a number of samples (*n* = 61) yielded an average number of spermatids per bundle of 63.15*±* 1.31, suggesting that developing pre-meiotic male germline cysts do in fact contain, on average, 16 cells.

## Acknowledgments

This study was supported by the National Institutes of Health (F31 HD098835 to R.D., R01 GM134204 to S.Y.S.) and the Howard Hughes Medical Institute (Y.M.Y.). The authors thank Tomer Stern for coding expertise, Sasha Meshinchi and the University of Michigan Microscopy Core for help with EM experiments, Paul Schedl for fly lines and antibodies, and Trudi Schüpbach, Eric Wieschaus, Paul Schedl, Hayden Nunley, Matt Smart, and Tatyana Gavrilchenko for helpful discussions and feedback.

## Notes

### Competing Interest Statement

The authors have declared no competing interest.

### Summary of Updates

The previous version of the manuscript focused only on the spermatogenesis work. The updated manuscript now includes methods for reconstruction volumes in female cysts, as well as a model analyzing the volume progression during Drosophila oogenesis.

