## Supplementary figures and images for "Fusome topology and inheritance during insect gametogenesis"

### Supplemental Figure 1

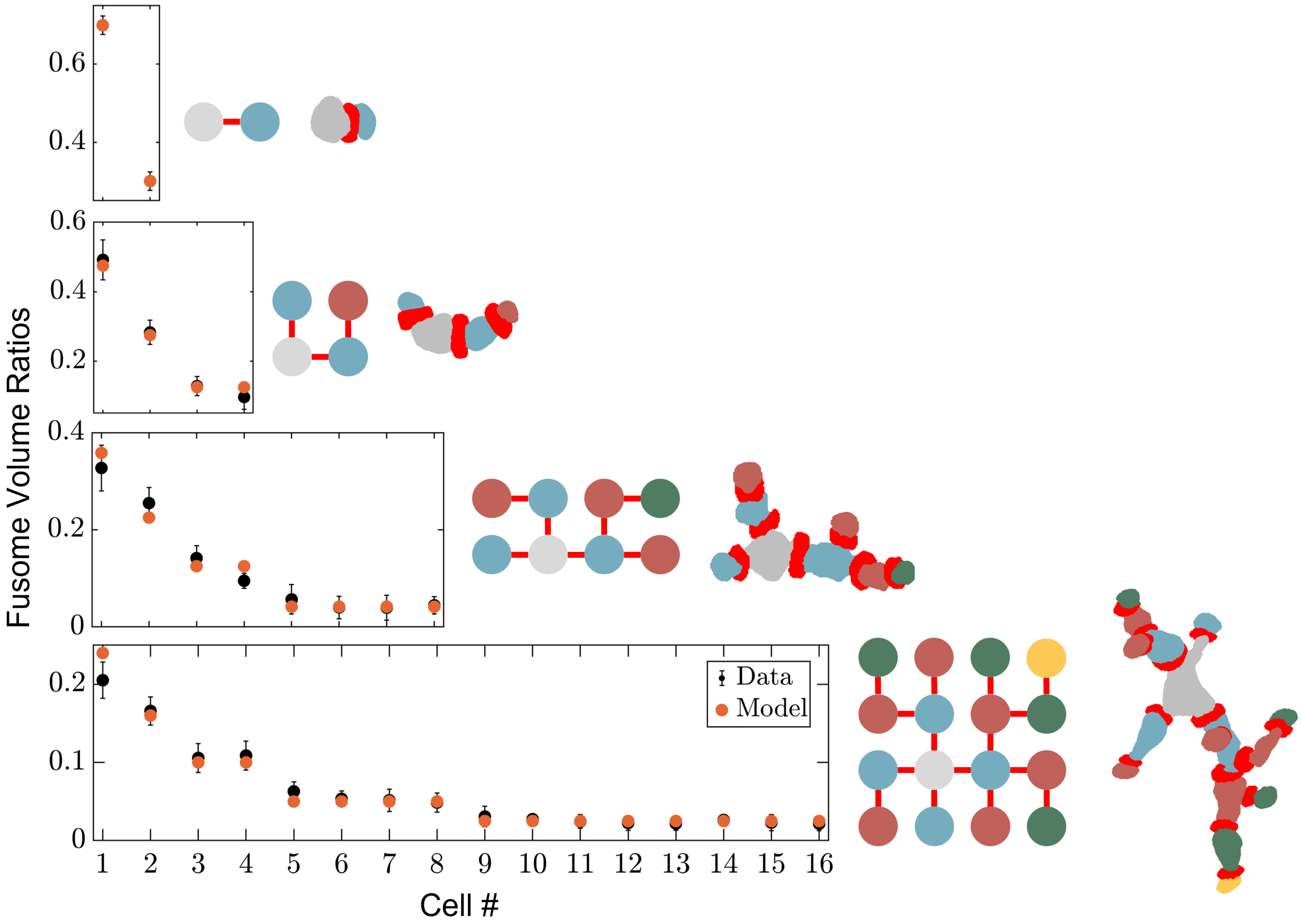
